# Implied motion guides neither gaze following nor intention attribution

**DOI:** 10.1101/2024.10.15.618221

**Authors:** Marius Görner, Masih Shafiei, Peter Thier

**Author notes:** Corresponding Authors: Marius Görner, Max Planck Institute of Psychiatry, Department Emotion Research, 80804 Munich, Germany. Masih Shafiei, Hoppe-Seyler-Str. 27, 72076 Tübingen, Germany. Peter Thier, Hoppe-Seyler-Str. 27, 72076 Tübingen, Germany. M.G. and M.S. contributed equally to this work. **Author Contributions:** M.G., M.S., and P.T. contributed to conceptualization, methodology, and writing – review & editing; M.G. and M.S. contributed to validation, visualization, and writing – original draft; M.G. contributed to data curation and formal analysis; M.S. contributed to investigation and Publication; P.T. contributed to funding acquisition, resources, and supervision. **Author Approvals:** all authors have seen and approved the manuscript, and that it hasn’t been accepted or published elsewhere.

## Abstract

The “gaze beam hypothesis (GBH)” of gaze following posits that the other’s eyes emit imaginary beams of moving energy travelling to the other’s object of attention (1), drawing the observer’s attention to the same object. This idea was initially supported by behavioral experiments showing a motion aftereffect (MAE), indicated by longer reaction times in detecting motion direction after viewing a cartoon face looking at an object in the same direction (2). However, this effect could also be expected if the observer used gaze direction to assume an intentional link between the looker and the object, envisioning directed actions toward the latter (3). To critically compare the two hypotheses, we tested whether an MAE could be induced by having human subjects detect motion direction after viewing various cue images, designed to differentiate the explanatory power of the two. Cues either suggested a connection between an agent and an object through the agent’s gaze or an object-oriented intention by the presence of equipment in the agent’s hand, without the agent directly looking at the object. Using Bayesian statistics, our findings provided strong evidence against both hypotheses at the population level, as reaction time modulations did not align with the MAE, leading us to reject motion adaptation as the underlying mechanism for gaze following but also intention attribution. As on an individual level we observed highly diverse effects, with some compatible with one or the other hypothesis, we assume that individual subjects may resort to different perceptual strategies based on different scene interpretations.

## Introduction

Viable social interaction requires the identification of the interests of others. One route to acquiring the needed knowledge is the other’s gaze, directed at objects of interest. It allows us, for our part, to shift our attentional focus to the same object, thereby establishing joint attention. But how we might link the other’s gaze to the object of interest remains unknown. An intriguing explanation was recently put forward by Guterstam et al. (1–4) who suggested that the subliminal perceptual experience of a force carrying beam emanating from the other’s eyes travelling to the object of interest might draw the observer’s attentional focus to the object, exploiting the implied motion associated with the putative moving beam (1, 2, 4, 5). A key finding supporting the idea of a force carrying beam, reported in a first publication (1) was that an observer perceived the other’s gaze to stabilize a vertical object, increasingly tilting towards the other. This was indicated by measuring that the tilt angle at which the observer expected the object to fall over was larger towards the side of the person gazing at the object (1). In subsequent studies, they presented evidence that the assumed force carrying beam had indeed motion-like properties (2, 4). This conclusion was suggested by the fact that observing a static image of a face gazing at a tree (*face* stimulus, Fig. 1) influenced reaction times (RT) when reporting the motion direction of a subsequently presented random dot motion stimulus (RDMS) (2). RTs in the incongruent condition, in which the direction of the RDMS was opposite to the gaze direction in the preceding cue image, were faster than in the congruent condition (*congruent deceleration*) (2). A similar differential effect on RTs characterizes the standard motion adaptation effect (6–8), in which an adapting motion pattern precedes judgments on motion direction. In fact, motion adaptation may also be induced by static images lacking real motion but implying motion such as a photograph of a running animal (7, 8). Against this backdrop Guterstam et al. concluded that also the other’s gaze, depicted in the static image directed towards the tree would imply motion and consequently induce motion adaptation. We refer to this idea as the gaze-beam hypothesis or GBH. The neural underpinning of this concept was investigated in an fMRI study in which Guterstam and colleagues trained a classifier to discriminate motion direction based on BOLD activity recorded when the participants were viewing coherently moving dots (4). Using the trained classifier, they then found higher classification accuracy in cortical areas known to be involved in visual motion processing (MT+) when the observer viewed a cartoon face gazing at an object compared to when the cartoon face was blindfolded (4).

**Figure 1.**
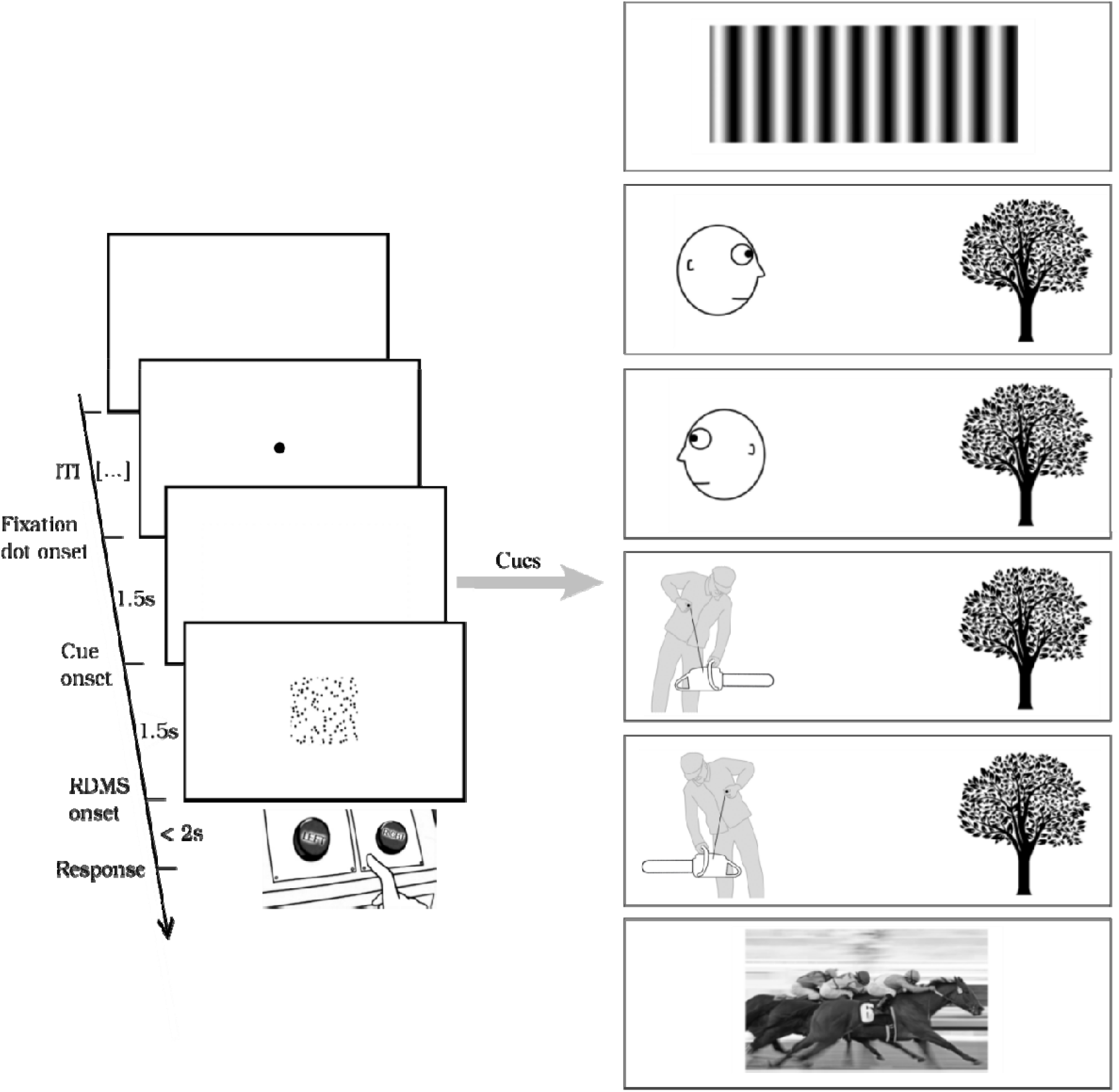
Illustration of the paradigm. The sequence of events in each main trial is depicted on the left side. Each trial started with a central fixation dot displayed for 1.5 seconds, following an inter-trial interval (ITI) of 250-500 ms. Next, one of the cues shown on the right side appeared on the screen for another 1.5 seconds. The cue was then replaced by a random dot motion stimulus (RDMS). Participants were instructed to quickly and accurately determine the direction of motion—either rightward or leftward—by pressing the corresponding key on the button box. Responses had to be made within a 2-s window. From top to bottom, the conditions are: *grating, face, mirrored*-*face, PWC, mirrored*-*PWC*, and *horses*.

In a commentary, Görner and colleagues argued against the GBH by pointing out its conceptual limitations and suggested an alternative explanation of the findings reported (3). They proposed that the observer’s expectation of motion of the agent and/or the object depicted in the image could be the cause of the activation of MT+. They reasoned that the observer will inevitably perceive the agent in the image as an intentional being about to manipulate the object, associated with the expectation of motion (9, 10). Such predictions of the other s intention are needed in order to prepare proper actions and, more generally, to shape the observer’s own behavior which is fundamental to successful social interactions (11). In fact, the idea that the expectation of movement may activate visual motion processing areas is well supported by previous work on the effects of imagined, predicted and remembered motion (9, 10, 12–14). Here, we refer to the idea that the attribution of intentions may evoke percepts of motion as the intention-attribution hypothesis (IAH). Importantly, unlike the GBH, the IAH predicts that assigning intentions to others recruits MT+ if the semantics of the scene suggest an object directed intention even though the agent depicted in an image does not yet attend the object.

In the present preregistered (15) study, we first repeated the original experiment of Guterstam and colleagues (2) using the two main cue images, a face looking towards a tree or, alternatively, gazing in the opposite direction (*mirrored-face* cue, Fig. 1). Guterstam and colleagues had previously reported that the *mirrored-face* cue did not result in a change in reaction time (no *congruent deceleration*). Secondly, we tested the IAH by investigating whether the motion adaptation effect is elicited by the rendition of a scene that implies object-oriented intentions of a person. This was implemented through an image that features a forest-worker standing a few steps away from a tree and pulling the starter rope of a chainsaw (*person-with-chainsaw* or *PWC*; Fig. 1). A key feature is that the worker’s eyes are obscured behind the visor of a cap and that the head is oriented downwards towards the chainsaw rather than to the tree, while the chainsaw is pointing to the tree. We assume that the compelling interpretation of this image is that the person intends to move towards the tree to fell it while his immediate attention is directed towards the chainsaw. Thereby, the *PWC* condition allows to distinguish intention (to cut the tree) from visual attention (devoted to the chainsaw). To test whether the orientation of the chainsaw matters, we included a mirrored version of the *PWC* cue image, in which the tool points to the opposite side. Further, as Guterstam et al. proposed that the gaze-effect is based on perceiving the cue image as implied motion (2, 4), we added a dedicated control of implied motion to the experiment by resorting to a static image of horses running to the left or right. Lastly, to test the ability of our experimental setup to reveal the motion after-effect elicited by real motion, we recreated the moving grating stimulus used by Guterstam et al. in their original study (2).

The hypotheses specified in the preregistered research plan (15) (see Table S1) were tested for each participant individually (Table S1, Q1-6) as well as for the population (ibid., Q7&8) (see *Transparent Changes*). To ensure the reliability of our results, we used Bayesian analysis methods and continued data collection from each participant until the calculated Bayes Factors met the threshold of *strong* evidence or until the number of trials per condition reached a maximum number in favor of the respective null/alternative hypothesis (see Methods for more details). For the population analysis, we proceeded with adding participants to our sample until the same evidence criterion was met.

Our results contradict Guterstam’s earlier results (GBH) but also the alternative hypothesis proposed by Görner et al. (16). On the population level we found *strong* evidence in favor of the null hypotheses that the cue images have no effect on subsequent judgments of the motion direction of RDM stimuli. However, the grating stimulus, as the control condition, elicited strong motion adaptation effects, validating our experimental setup. Analysis at participant level revealed a surprisingly diverse picture. Although the majority of participants still showed a null effect in most conditions, we did find individuals whose data suggested highly idiosyncratic effects with *strong* evidence. Some of them were notably slower in detecting the dot motion direction when it was congruent with the direction depicted in the cue image, whereas others were faster. This finding highlights the importance of participant-level analyses in general. At a conceptual level, it shows that the effects reported by Guterstam et al. (1, 2, 4, 5) and here may arguably reflect individually different experiences and subjective interpretations of the scenes depicted in the stimuli.

## Methods

### Transparent changes

In deviation from the original research plan (15) we based the population level analysis not on the distribution of individual effect directions but on population level estimates of effects. This is in greater alignment with analysis methods employed by Guterstam et al.

### Ethics information

The research presented here was approved by the Ethics Board of the responsible institution and was conducted in accordance with the principles of human research ethics of the Declaration of Helsinki. Informed consent was obtained from all participants, and they were compensated with 7€ per half hour.

### Design

Our study consisted of two parts, a pre-experiment, and the main experiment. Both involved motion-discrimination reaction-time tasks. Researchers were blind to the experimental conditions during data collection. Analysis was not blinded.

The pre-experiment consisted of 100 trials. In each trial, participants first saw a red central fixation dot (0.5° in diameter) which was shortly after replaced by a random dot motion stimulus (RDMS) with varying noise levels. The movement direction of each individual dot was sampled from a von Mises distribution with a mean of either 90° or 180° (rightward or leftward motion, randomized across trials) and varying precision (noise levels). The dots had a lifetime of 200 ms and a velocity of 2 °/s. The RDMS was presented in a 5-by-5° square at the center of the screen with black dots on a white background. This procedure generated a flow field with a dominant overall direction of motion. Participants were instructed to report the direction as quickly and accurately as possible by pressing the respective button on a button box. Goals of the pre-experiment were to familiarize participants with the discrimination task, and to find the individual noise level at which the participants’ psychometric function reached an 80% correct response rate. Our pilot data reported in the registered research plan (15) suggested that the motion adaptation effect was strongest at this noise level.

Our main experiment followed a within-subject design. Participants carried out the same motion discrimination task using the RDMS as in the pre-experiment, with two modifications: first, the noise level was tailored to each individual’s performance as measured in the pre-experiment, and second, cue images preceded the RDMS. Figure 1 illustrates the temporal sequence of a main trial as well as the different cue images. The cues were either a sinusoidal gray scale moving grating or one out of five different sets of images (referred to as the *cue conditions*). The grating had a period of 0.8° and measured 14.7° in width by 5.7° in height. It moved to either left or right at a speed of 0.8°/s for 1.5s, while its rectangular boundary remained immobile. In four of these sets, an image of a tree was presented on one side of the screen, right or left with equal probability, while on the other side, one of the following images appeared: *face, mirrored*-*face, PWC* and *mirrored-PWC*. Mirrored means that the *face*/*PWC* were facing away from the tree. In the fifth one, a single static image of horses, running to either right or left selected at random, was presented centrally. The distance between the two images was equal to the width of the RDMS (≈5°). Dimensions of the *PWC* were chosen to match those of the *face* image. After a variable inter-trial interval (ITI) of 250 – 500 ms, the central fixation dot (0.5° in diameter) was presented for 1.5 s, followed by the cue shown for 1.5 s. As soon as the RDMS appeared, participants had 2s to report its direction of motion. Participants sat in front of the display in a dimly lit room, resting their heads on a chinrest to help maintain a 103 cm distance from the screen (597 mm by 340 mm). They were instructed to keep their gaze near the screen’s center (where the fixation dot had been located) during the cue period and to avoid making exploratory saccades to scrutinize the images. To familiarize them with the cue images and reduce the urge to closely examine them, all cues were shown to the participants before transitioning to the main experiment. The experimenter provided a brief description of each cue image, such as “a face is on the right and a tree on the left,” or “a person holding a chainsaw is on the left and a tree on the right,” to ensure they understood the image contents. Moreover, they were told to press the button corresponding to the perceived motion direction as quickly and as accurately as possible once the RDMS appeared on the screen. Upon responding, the RDMS disappeared, signaling the start of ITI. Participants had a maximum of 2 seconds to indicate their direction choice; if they failed to do so, the RDMS vanished, and a “too slow!” message was displayed for 5 seconds. The ITI began after this message was cleared from the screen. Any trials labeled “too slow” were repeated at the end of the same run.

Four the four cue images involving a tree and a person, the position of the tree on the left or the right side of the image served as the reference direction for the comparison with the direction of the RDMS, independent of the orientation of the face or the assumed direction of intention. For the remaining two stimuli, the direction in which the horses were running and the direction of movement of the grating were used as reference direction. When the cue’s reference direction matched the motion direction in the subsequent RDMS, the trial was considered congruent; otherwise, it was incongruent.

The layout of the *face* images, the moving grating, and RDMS as well as the temporal sequence and duration of presentation of stimuli, were adopted from Guterstam *et al*. (2). We deviated from Guterstam et al. in the algorithm with which we generated the RDMS. We drew the motion directions of individual dots from a von Mises distribution while they used a subset of dots moving in the respective direction with all others moving in random directions. Our method has the advantage that it prohibits the strategy to use individual dots to identify the motion direction. Further, Guterstam et al. used a fixed noise level (i.e. the percentage of dots moving in random directions) for all participants and excluded those who did not achieve at least 80 % accuracy. We, on the other hand, adapted the noise level to each participant’s individual capabilities, using the 80% accuracy as the threshold.

Data for the main trials was collected over several days, with each session lasting up to 90 minutes. In each session, trials were organized into runs of 50-60 trials, featuring a maximum of three different *cue-conditions*. Daily sessions could include multiple runs from a subset of conditions. Within each run, trials of each cue category and the two congruency conditions (congruent vs. incongruent configurations) were randomly interleaved and counterbalanced. To eliminate the influence of the motion aftereffect elicited by the moving grating on the performance in other cue conditions in subsequent trials, the moving grating was tested in separate runs. We limited the number of conditions within each run to minimize the likelihood of outliers in *RT* as we included *run* as a group-level effect in the regression model. Participants were permitted to take a short break between runs upon request. Participants performed the mirrored conditions of the *face* and *PWC* stimuli only if they showed a difference between the RTs of congruent and incongruent conditions in the non-mirrored variants.

### Data collection

We used a Bayesian updating approach for data collection. According to the guidelines for registered reports that employ Bayesian hypothesis testing (17), we aimed at collecting data until the Bayes Factors (*BFs) were at l*east 10 in favor of the null or the alternative hypotheses at both the participant and population levels. The first part of the analysis focused on the participant level reaction times (RTs), analyzing each participant’s data individually for differences between the reaction times of congruent (*RT*_*c*_) and incongruent (*RT*_*ic*_) trials. The second part addressed the population-level effects. We continued collecting data from each participant and added further participants to the sample until the BF criterion was reached. This was specifically important for the main control condition (*Grating*) and the two main experimental conditions (*Face* and *PWC*). Reliable results in the other conditions (*mirrored-face, mirrored-PWC and Horses*) were only required as additional controls in case of an effect in the two main experimental conditions. For practical reasons, we needed an additional stopping rule for data collection from individual participants. This involved setting a maximum number of runs per condition, which was 25. Initially, 11 participants were enrolled in the study. After applying these inclusion criteria, data from 9 participants with varying trial numbers were included in the final analysis (see Table 1). The remaining 2 participants were unable to continue for personal reasons unrelated to the study, resulting in incomplete datasets that were excluded from the final analysis.

**Table 1.** Bayes Factors (0/1*) of the po*pulation level models.

|  | *grating* | *face* | *PWC* | *horses* | *mirrored-face* | *mirrored-PWC* |
| --- | --- | --- | --- | --- | --- | --- |
| $BF_{01}$ | -4.3e-33 | 11.0 | 13.5 | 7.11 | 2.6 | 10.6 |
| $\Delta RT$ (ms) | 71.2 | -4.7 | -4.3 | 9.2 | 8.4 | -4.4 |
| $CI^{95\%}$(ms) | [50.3, 91.3] | [-14.1, 9.2] | [-12.9, 4.2] | [-4.7, 18.2] | [0.0, 16.6] | [-18.0, 8.7] |

Note that positive values indicate shorter RTs in the incongruent condition, consistent with the expected effect of *congruent deceleration*.
$\Delta RT$ = mean $RT_c$ – mean $RT_{ic}$ ; $CI^{95\%}$ : 95% credible interval.

### Analysis

The analysis was based on trials with correct responses only. No further outlier removal was performed. All analyses were conducted using custom-made scripts in MATLAB (18), and R (19). For Bayesian modeling and hypothesis testing, the R package *brms* (20) was used.

### Differences in reaction time and its variance at the participant level

We were interested in the effect of the congruence-condition in which the directions of the cues matched the direction of the subsequent dot motion, grouped by cue-condition, on RTs. RTs were analyzed for each participant individually. To this end, for each cue-condition we fitted (distributional) Bayesian hierarchical models with the congruency-condition as the population-level (fixed) effect and runs as group-level (random) predictor with random intercepts and slopes (see Supplementary material for the detailed model specification). We used the congruence-condition not only as a predictor for the mean but also for the variance, since pilot data suggested that effects on the variance existed as well. The outcome variable was truncated at an upper bound of 2000 ms reflecting the maximum RTs allowed by the experimental design. *Runs* were only included as group-level factors if the number of runs was larger than 5 as suggested by Bolker (21). Priors for the outcome variables are tuned such that prior and posterior predictive checks yield reasonable results for the pilot data and simulated data (see Supplementary Material - Model specification and the registered research plan (15) for details). The priors for the effects of interest were chosen to be weakly informative, reflecting our expectation that effects on RTs are within a range of ±250 ms which is the range where this prior has roughly 80% of its mass. We computed *BFs for the poi*nt-Null-hypotheses specified in Table S1 and described in more detail in the registered research plan (15) using the Savage-Dickey (22) approach which computes *BFs as the rati*o between the prior and the posterior distributions at the specified parameter value, here 0. This procedure is equivalent to a two-sided test. The model was run using the standard settings of *brms* (4 Markov chains with 2000 iterations, each). In addition to the *BFs, we report* the estimates for *Δμ*_*RT*_ and their 95% Credible Intervals. For intuitive understanding, the estimates for *Δμ*_*RT*_ will be transformed into *ms* using the formula 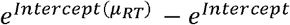 because internally the model is on the *log scale. Thro*ughout the manuscript we report Bayes Factors in the form 0/1 which means that *BFs* > 0 *represent e*vidence in favor of the *null*-hypothesis and *BFs* < 0 *evidence in* favor of the *alternative*.

### Testing the validity of the GBH and the IAH at the population level

To test whether any of the two hypotheses is supported at the population level, we computed Bayesian hierarchical models with the congruence-condition as the population-level (fixed) effect and *participants* with nested *runs* as group-level (random) effects with random intercepts and slopes across conditions (see Supplementary material for the model specification). As prior for the main effect we used the same one as above and Bayes Factors were computed in the same way as described above, as well. The *GBH* and the *IAH* both require a specific pattern of effects across the conditions in order to be supported (see Table 1, Q7 & Q8). In short, *GBH* predicts that, aside from the control conditions, there is a motion-adaptation effect only for the *Face* condition and no effect for the *mirrored-face, PWC* and *mirrored-PWC* conditions. IAH predicts that the *PWC* and *mirrored-PWC* conditions show motion adaptation effects as well.

## Results

### Participant level

Figure 2 summarizes the participant-level results. We found *strong* evidence (*BF* > 10) *for the* “congruent deceleration” effect (*RT*_*c*_ > *RT*_*ic*_) in the grating condition for all participants except two (P3 and P7). This finding validated that our behavioral apparatus could capture the “motion aftereffect”. Participant 7 was unable to complete data collection for this condition. In all other conditions, the majority of the participants (five to seven out of nine) showed moderate to strong evidence supporting the null effect (no difference between *RT*_*c*_ and *RT*_*ic*_). Among the remaining participants, some demonstrated “congruent acceleration” (*RT*_*c*_ < *RT*_*ic*_) in certain cue conditions, while others displayed mixed effects. For example, participant 3 showed “congruent acceleration” in the *face, Horses* and *PWC* conditions, whereas participant 9 presented “congruent deceleration” in *face, mirrored-face*, and *Horses* conditions, but the opposite effect in the *mirrored-PWC*. Only one other participant (P1) exhibited “congruent deceleration” in the *Face* condition, and none of the participants showed “congruent deceleration” in the *PWC* or *mirrored*-*PWC* conditions. Furthermore, two participants (P8 and P9) did not provide sufficient evidence even the maximum amount of data collected (25 runs) in the *Face* and *mirrored-face* conditions (P8) or the *PWC* condition (P9). Overall, participant-level analysis revealed no consistent effect pattern within or between participants that would correspond to the respective predictions of the *GBH* and *IAH*. Supplementary Table 1 provides a detailed overview of the individual participant-level results.

**Figure 2.**
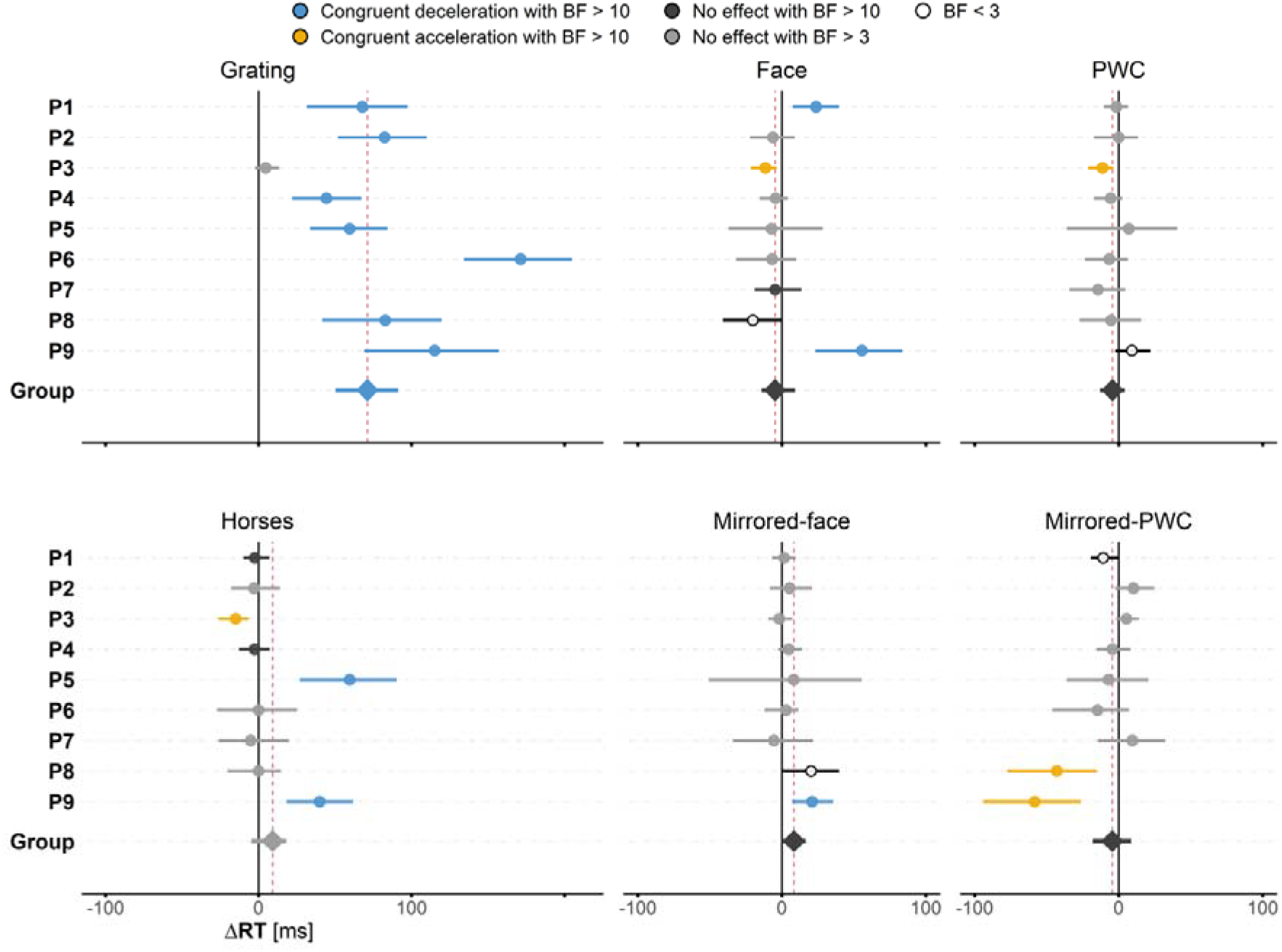
Summary of *ΔRT at both the* participant-level (dots) and group-level (diamonds), calculated by subtracting the mean *RT*_*ic*_ from the mean *RT*_*c*_ results (see supplementary Table 1 for more details). Positive values indicate shorter mean *RT*_*ic*_, suggesting “congruent deceleration” or “motion adaptation.” In contrast, negative values correspond to shorter *RT*_*c*_, reflecting “congruent acceleration”. Horizontal bars denote the 95% credible interval (*CI*^95%^), while vertical solid lines mark the zero point, indicating no change in RTs. Vertical dashed lines represent the group-level estimate. Colors and shades of gray depict the direction of effect and the strength of evidence (see figure legend). Open circles (*BF* < 3) mark data points with insufficient evidence. Note that participant 7 did not complete data collection for the *grating* condition.

### Group level

Figure 2 and Table 2 summarize the Bayes Factors, effect estimates and *CI*^95%^ of the population level analysis. The only condition for which we found evidence in favor of an effect of the cue on RTs at the population level is the *Grating* condition, reaffirming the validity of our setup. In both main experimental conditions (*Face* and *PWC*), our data provided *strong* evidence against any effects, falsifying both *GBH* and *IAH* at the population level. Also, for the *Horses* and the *mirrored-PWC* conditions, our data provided *moderate* and *strong* evidence against any effect, respectively. With a *BF* < 3, *the data* of the *mirrored-face* condition provided insufficient evidence to draw any conclusions which is, however, irrelevant for interpretation considering the outcomes of the other conditions.

## Discussion

In the present study, we attempted to compare the explanatory power of two competing hypotheses proposed to explain the motion adaptation effect evoked by object-directed gaze. The first one was formulated by Guterstam et al., suggesting that the observer’s brain perceives other’s gaze as imaginary force carrying beams moving from the eyes of the looker to the target refer to as the gaze beam hypothesis (GBH) in this article. According to Guterstam et al. (1), the assumed motion character of the beams was established by demonstrating that object-directed gaze gives rise to a motion aftereffect (MAE). The alternative hypothesis, termed the intention attribution hypothesis (IAH), proposed by us, assumes the emergence of an intentional bond between the other and the object she/he is attending to(3). As intention attribution entails the expectation that the other may launch an action on the object necessarily involving directed motion, the observed MAE might in fact be secondary to intention attribution. As previous experiments were not designed to compare the viability of the two contrasting hypotheses, we here set up experiments able to discriminate between the two. To enhance transparency and credibility, and to reduce the risk of a biased reporting, we pre-registered the present study, detailing the conceptual framework of the GBH and the IAH and the criteria for accepting or rejecting each hypothesis. An important aspect of the study plan was a clear roadmap for the collection and analysis of data resorting to Bayesian statistics. In order to reevaluate the evidence for motion adaptation as indicator of implied motion associated with the force carrying beam assumed by the GBH, we replicated Guterstam and colleagues’ experiments that suggested motion adaption as a consequence of a preceding object directed gaze stimulus (1). However, the data obtained by us could not corroborate this result. In fact, at the population-level the data provided strong evidence in favor of the null-hypothesis – no motion aftereffect following gaze stimuli, thus contradicting the predictions of the GBH– although our subjects exhibited clear motion aftereffects when exposed to standard motion stimuli, verifying our experimental setup. In order to critically assess the viability of the alternative, the IAH, our experiment resorted to new stimuli, tailored to dissociate the two hypotheses. The central idea was to present an actor, seen to prepare an object directed action, yet without offering gaze cues. However, also here the population data contradicted the predictions of the IAH with strong evidence. Considering the results from both parts of our experiment – those aiming at the GBH and those at the IAH – we were forced to reject both. However, this conclusion does not rule out that individual subjects may have adopted idiosyncratic perceptual strategies, compatible with one or the other hypothesis, but not necessarily stable across the participants, a possible interpretation that will be discussed in more detail below.

Both the GBH and the IAH require that subjects are able to experience implied motion, able to modulate the perception of real motion, no matter if the implied motion stimulus is a force carrying beam emanating from the eyes or a directed intention. Hence, one might argue that our failure to obtain results consistent with either hypothesis might be secondary to a fundamental lack of sensitivity to implied motion. This is why our experiment involved a component allowing to assess our subjects’ ability to experience a MAE elicited by a more conventional implied motion stimulus, namely a static image of a group of horses stampeding in a particular direction. Finally, subjects were also tested in a control condition in which a *moving grating* stimulus was used to evoke a MAE. Consistent with Guterstam et al.’s findings (1), our results demonstrated a MAE associated with the *grating* stimulus, with moderate to strong evidence at both the individual participant level and strong evidence at the population level. This confirms the effectiveness of our behavioral paradigm in capturing MAEs and suggests that the lack of a MAE in the *face* and *PWC* conditions across participants, which led to the rejection of GBH or IAH, cannot be attributed to a faulty experimental setup.

Unlike the *grating* stimulus, an “implied motion” stimulus, such as *horses*, was not included in any study by Guterstam et al. (1, 2, 4, 5). Our results indicated moderate evidence in support of a null effect in the *horses* condition, suggesting the possibility that the paradigm may not have effectively evoked implied motion evidenced by a MAE. While this finding raises the concern that the setup used in this study and the one reported by Guterstam and colleagues may be unable to unravel an impact of implied motion on the perception of real motion, it fails to explain the discrepancy between our inability to reproduce the positive evidence in favor of the GBH presented by Guterstam et al., given the fact that the stimuli between their and our experiments were identical. Possible explanations for the inability of the *horses* stimulus to elicit MAE include the extended duration of its presentation (1.5 seconds) and the repeated use of the same static image throughout the experiment. In contrast, most previous studies that successfully triggered the MAE with implied motion stimulus used sequences of different images, shuffled before each presentation (35-37). Each image was typically shown for about a third of the time used in our experiment – around 500 milliseconds – while the entire sequence would last anywhere from a few seconds to several tens of seconds. It seems likely that longer presentation duration of the same stimulus favors the perceptual interpretation that the scene is actually motionless. Our choice of the implied motion stimulus and presentation parameters were made with the aim of ensuring that the stimulus had characteristics comparable to the *face* and *PWC* stimuli, allowing for meaningful comparisons across different cue conditions. Future research could focus on structuring a paradigm for reliably inducing and measuring implied motion using widely accepted stimuli, complementing stimuli related to gaze following and intention attribution.

Our experimental design enabled us to delve into participant-level data, revealing insights that would have been obscured if we had relied solely on population-level results. Indeed, by examining individual data, we uncovered a significant diversity in the effects observed across participants. For instance, participant 1 and 9 demonstrated a congruent deceleration effect in the *face* condition with strong evidence, partially supporting the GBH. However, their overall results did not meet all the criteria necessary to fully endorse the GBH. Specifically, participant 9 showed strong evidence in favor of a congruent deceleration effect in the *mirrored-face* condition, while participant 1 presented strong evidence for a null effect in the *horses* condition, both findings that contradict the GBH predictions (which required null effect in the *mirrored-face* condition and congruent deceleration in the *horses* condition). Furthermore, our analysis revealed a congruent acceleration effect with strong evidence across different stimuli in other participants (e.g., P3, P8, and P9). These variabilities in effect types, both within and between participants, point to individually different perceptual strategies deployed when trying to interpret a scene. Moreover, these strategies might also be condition dependent. This is indicated by the fact that an individual may present a congruent acceleration effect in one condition (e.g. subject P8), a congruent deceleration in another and null effects in yet others. The alternative possibility that these effects are due to chance does not seem plausible given the fact that the Bayes Factors indicated strong evidence for each. The condition dependency of putative perceptual strategies might be due to a shift of focus on particular visual features in the scene, depending on the interpretation it prompts. To avoid different interpretations of the cues, future research could include a pre-cue stimulus priming participants’ interpretation, ensuring a more consistent understanding of the cues across participants and therefore allowing for a clearer assessment of its impact on motion direction discrimination.

One intriguing participant-level effect observed was the congruent acceleration (shorter RTs in the congruent rather than the incongruent condition), occurring for some participants across a few stimuli. These stimuli (such as horses, faces, etc.) appeared to have no clear similarities, aside from potentially offering directional information. These stimuli (such as *horses, face*, etc.) did not share any obvious similarities apart from providing directional information. We speculate that these disparate acceleration effects may be driven by feature-based attention to direction (28). Attention to direction may then increase the sensitivity to the same direction in the subsequent RDM test pattern enhancing the detection of motion in this direction. A faciliatory impact of attention to motion direction is well-established and probably based on the properties of neurons in motion sensitive cortex MT+ visual areas (25, 27, 28). Alternatively, the congruent acceleration might be explained by response priming. Here the directional cue triggers motor preparation, thereby speeding up the execution of the required action. Unlike the GBH, IAH, and feature-based attention, which attribute the cue effect to a modulation of neuronal activity in the visual motion complex MT+, response priming assumes that the cue-induced modulation occurs primarily at the motor-level. The cued direction triggers the preparation of the motor response that has to be executed if the test stimulus’s direction matches the cued direction. This interpretation aligns with previous research suggesting that the classic “Posner spatial cueing effect” might be more accurately explained by a response priming mechanism within the motor domain rather than by spatial attention allocated to the position of the target stimulus. Our paradigm can be compared to the “Posner cueing paradigm,” as both share a key element: the influence of a task-irrelevant cue on subsequent visual detection.

The GBH has gained support from an fMRI study in which whole-brain BOLD activity was recorded from human subjects as they viewed three different stimuli: coherently moving dots (rightward or leftward) or a cartoon human head facing rightward or leftward at an object, with eyes either open or blindfolded. A classifier that had learned to differentiate motion direction based on MT+ BOLD activity evoked by the moving dots, distinguished rightward versus leftward gaze more accurately in the eyes-open than the eyes-covered, in right MT+ voxels preselected for their preference for the eyes-open. This finding was interpreted as supporting the assumption that motion perception arises from observing open eyes gazing at an object, a percept that is no longer evoked if the eyes are covered and therefore in line with the basic premise of the GBH. However, the very study also showed that the classifier’s ability to detect object-directed gaze, was not entirely absent, although significantly less accurate in the eyes-closed condition, when only voxels preferring eyes-open were considered. In other words, the reported MT+ activity is not confined to gaze-based motion perception, a conclusion that aligns with the studies showing that this region is also engaged when attention is directed to objects via cues like arrows (30-32). In fact, a previous human fMRI study demonstrated that MT+ activity does not distinguish between an arrow or a gaze stimulus guiding attention to an object of potential interest (32). This is of special interest here because in their original behavioral study (1), Guterstam et al. did not find any behavioral effect of arrow stimuli. Hence these findings imply that MT+ activity might reflect a broader role in processing spatial and feature-based attention. This role extends beyond motion-related processing, explicitly driven by moving objects. Additional evidence for this broader function comes from human imaging studies linking modulation of MT+ activity with auditory attention (33) and neurophysiological studies in nonhuman primates indicating an MT+ involvement in feature-based attention, such as to color (34), as well as the processing of low-level visual features (38).

In an earlier study, Guterstam and colleagues demonstrated that observing a cartoon face looking toward a vertical bar influenced participants’ judgments of the tilt angle at which the bar was perceived as falling over was larger when tilting toward the face compared to when the tilting was away from the face. The authors interpretated the observed stabilizing effect as if imaginary force carrying beams were extended from the cartoon face toward the vertical bar. Moreover, they suggested that the experience of this beam by the observers was associated with the perception of directed movement. Considering the evidence against the experience of gaze associated motion presented here, one may wonder if there is an alternative interpretation of this intriguing behavioral finding? One plausible speculation might be that the observation of somebody visually attending to an object conveys the anticipation of an upcoming action, such as a hand reaching for the object, to provide support. This is consistent with the idea that object-directed visual attention often precedes coordinated hand movements. This concept closely parallels the Intention Attribution Hypothesis (IAH), which posits that perceptual consequences arise from implicitly inferring an intention to act based on visual attention. However, unlike the IAH, the suggested mechanism would not rely on implied motion. Instead, it may be grounded in broader Gestalt principles that explain causal relationships between agents and objects. This hypothesis could be further explored in experiments manipulating hand availability in the tilt judgment task.

In conclusion, despite replicating the methods and paradigm of Guterstam et al, we were unable to reproduce their findings supporting the GBH. Beyond that, we also failed to obtain evidence to support our alternative hypothesis, the IAH. These results cannot be dismissed as statistical noise, as our Bayesian analysis— which was carefully designed and transparently pre-registered prior to data collection—provided moderate to strong evidence against both hypotheses. This leads us to conclude that neither the GBH nor the IAH – two competing hypothesis that posit a role for implied motion associated with the gaze of the other – allow the observer to follow the other’s gaze to objects of interest.

## Acknowledgements

We thank Dr. Friedemann Bunjes and Peter W. Dicke for their invaluable technical assistance.

We acknowledge support from the German Research Foundation (DFG) project TH425/17-1 and from the Open Access Publication Fund of the University of Tübingen

